# Myofibroblast Forms Atherosclerotic Plaques

**DOI:** 10.1101/2020.07.20.212027

**Authors:** Xinggang Wang, Junbo Ge

**Author notes:** Corresponding author: Junbo Ge, 180 Fenglin Road, Shanghai, China 200032.

## Abstract

For decades, smooth muscle cells (SMCs) and macrophages are considered as the main contributors to atherosclerotic plaques. However, we found that in the human coronary atherosclerotic plaques, SMCs were few, while lots of myofibroblasts infiltrated in the intima near the lumen (fibrous cap) and their distribution was highly positive related to intimal thickness. In addition to lots of foam cells formation, collagen fibers were forming in the thickening intima near the lumen (fibrous cap), and denaturing or calcifying gradually far from the lumen, which evolved into various complex plaques. In vitro, myofibroblasts could actively take lots of low-density lipoprotein (LDL) to enhance proliferation. Lots of collagen fibers, foam cells and extracellular lipids accumulation emerged in myofibroblasts cultured with 5% FBS high glucose DMEM without adding modified LDL. It is consistent with the characteristics of human coronary atherosclerotic plaques. It is the first time that lipid rich plaques with lots of foam cells, extracellular lipids and collagen fibers formed in vitro. It demonstrated that myofibroblast should be the direct and main source of collagen fibers, foam cells and extracellular lipids. This suggests that atherosclerosis is not as complicated as previously considered, and it might be mainly a process of myofibroblast remodeling to vascular injury caused by various risk factors. This study made the pathogenesis of atherosclerosis clearer. It would provide a target cell for future treatments of atherosclerotic diseases. In vitro atherosclerotic plaques model formed by human myofibroblasts would be an efficient and convenient way to study atherosclerosis.

## Introduction

Atherosclerosis is the common reason of cardiovascular and cerebrovascular diseases of human beings^1^, while its pathogenesis is still unclear. It is a progressive disease characterized by foam cells formation, accumulation of extracellular lipids and fibrous tissues in the intima of the large and medium arteries^2^. It is now believed that it is the results of a combination of multiple risk factors, such as hypertension, hyperlipidemia, hyperglycemia, smoking, vasculitis, etc^3^. These risk factors could lead to vascular injury. The traditional views are that vascular injury, lipids deposition, macrophages and smooth muscle cells (SMCs) phagocytosis of lipids forming foam cells are the main contributors to atherosclerosis^2^. These studies made us to consider how repair and remodeling occur after vascular injury. Which cells are the direct performers to vascular repair and remodeling? Do they play an important role in the development of atherosclerosis? Fibrous tissue repair is a common repair way after severe injury of organs and tissues^4^. More and more studies found that fibrous tissues are widely distributed in atherosclerotic plaques^5–7^. Based on these previous studies, we speculated that atherogenesis might be a process of vascular fibrous tissues repair and remodeling to the vascular injury caused by various risk factors. Fibrous tissue remodeling might also be an important reason for the formation of foam cells and extracellular lipids accumulation, which have not been reported.

In humans, many cells could secrete extracellular matrix to form collagen fibers and participate in the fibrous tissue repair, such as SMCs, fibroblasts, myofibroblasts, etc^8^. In skin and other organs, some studies have found that when the tissue damage is serious or the mechanical force is significantly increased, fibroblast, SMCs or endothelial cells, etc. could transform into myofibroblast to perform its function of tissue repair and remodeling^9^. Therefore, compared with SMCs, endothelial cells and fibroblasts, myofibroblasts play a more important role in the fibrous tissue remodeling. Although during all stages of coronary lesions, including mildly stenotic plaques, highly stenotic stable plaques and erosions, and restenosis lesions, myofibroblasts are widely distributed in atherosclerotic plaques^9–11^, the relationships between fibrous tissue remodeling and foam cell formation has not been paid much attention. The roles of myofibroblast in atherosclerosis has been neglected. The traditional views are that macrophages and SMCs phagocytosis of lipids forming foam cells are the main contributors to atherosclerosis^2^. Recent studies reported SMCs are the main source of foam cells in human coronary artery, while macrophages derived foam cells account for only 3–5% of total cell populations^12, 13^. Benditt EP proposed that the SMCs in atherosclerotic plaques are monoclonal, a mutant SMC produces the progeny cells, migrates into the intima, divides and proliferates to form plaques, just like a leiomyoma^14, 15^. This hints that whether myofibroblast is the “mutant SMC” to play a key role in the repair and remodeling of coronary injury? Although myofibroblasts are similar to SMCs in morphology, there are many differences in proteins expression^16^. Alpha-smooth muscle actin (α-SMA) is highly expressed in SMCs and myofibroblasts, while smooth muscle myosin heavy chain (hc-SMM) is highly expressed in SMCs but lowly expressed in myofibroblasts^17^. In addition, myofibroblasts have stronger ability of proliferation and extracellular matrix secretion to perform tissue repair and remodeling compared with SMCs^10, 16^. Therefore, in order to study the role of myofibroblast in atherosclerosis, human coronary arteries were explored. Then we cultured myofibroblast in vitro. We found that myofibroblast formed lipid rich plaques with lots of foam cells, extracellular lipids and collagen fibers in vitro. Myofibroblast should be the direct performer of atherosclerosis, and fibrous tissue remodeling is an important reason of atherosclerosis.

## Materials and Methods

### Coronary artery specimens

Human coronary arteries (n=66) were collected from autopsies which conformed to the principles outlined in the Declaration of Helsinki and obtained written consent from their relatives prior to the inclusion of subjects in the study. They were fixed with formalin, washed with water, dehydrated with alcohol, embedded in paraffin, sectioned into 4 μm thick on the slides.

### HE Staining

Eosin and hematoxylin staining were used to the human coronary arteries with HE staining method.

### MASSON staining

The sections were dewaxed to water, stained with Masson Ponceau acid complex red solution, soaked with 2% glacial acetic acid solution, differentiated with 1% phosphomolybdic acid solution, stained with aniline blue, soaked with 0.2% glacial acetic acid solution for a while. Dehydrated in alcohol gradient, cleared in xylene, sealed with neutral gum, and photographed. Proportion of collagen fiber area to intima area was measured with Image-Pro Plus 6.0 and was analyzed with Origin 9.6.

### Immunofluorescence

The sections were dewaxed to water and repaired with hydrogen peroxide, washed with water, penetrated with Triton x-100, blocked with BSA for 2h, incubated with primary antibodies overnight at 4 ◻. Then they were washed 5min for 3 times with PBST. Fluorescence labeled secondary antibodies were added to the specimen for 2h in the dark, then washed for 5min for 3 times with PBST, and then sealed with glycerin and imaged. The data was measured with Image-Pro Plus 6.0 and was analyzed with Origin 9.6.

### Cells culture in vitro

WPMY-1 cell line provided by cells bank of Chinese Academy of Sciences. It is a frequently-used myofibroblast cell lines from human.

The cells were cultured in 24 wells plates in high glucose DMEM containing 5% FBS in incubator with 5% CO_2_ at 37 ◻. The cells were divided into two groups: the control (Con) group and the DiI-LDL group. The final concentration of DiI-LDL was 30 μg/ml, and the culture medium was sucked out and washed twice with PBS 24 hours later, then 5% FBS high glucose DMEM was added. The cells were then imaged under Leica DMi8. Each group was set with 3 multiple holes in each time. They were repeated for 3 times (n=9).

The cells labeled with DiI were cultured in 24 wells plates in the incubator with normal condition. At days 7, 14 and 21, they were imaged, stained with Oil Red O and Hematoxylin, and were imaged with Leica DMi8. Each group was set with 3 multiple holes in each time. They were repeated for 3 times (n=9).

### Statistical methods

The difference between two samples was tested by t test with origin 9.6, and P < was statistically significant. Linear fit and box plot were analyzed with origin 9.6.

## Results

### Myofibroblast was positively related with intimal thickening of human coronary artery

There were few SMCs in the thickening intima, while there were lots of myofibroblasts, particularly near the lumen (Fig.1A, 1B). Furthermore, myofibroblasts distribution is highly positive correlated with intimal thickness (Fig.1C,1D,1E).

**Fig.1.**
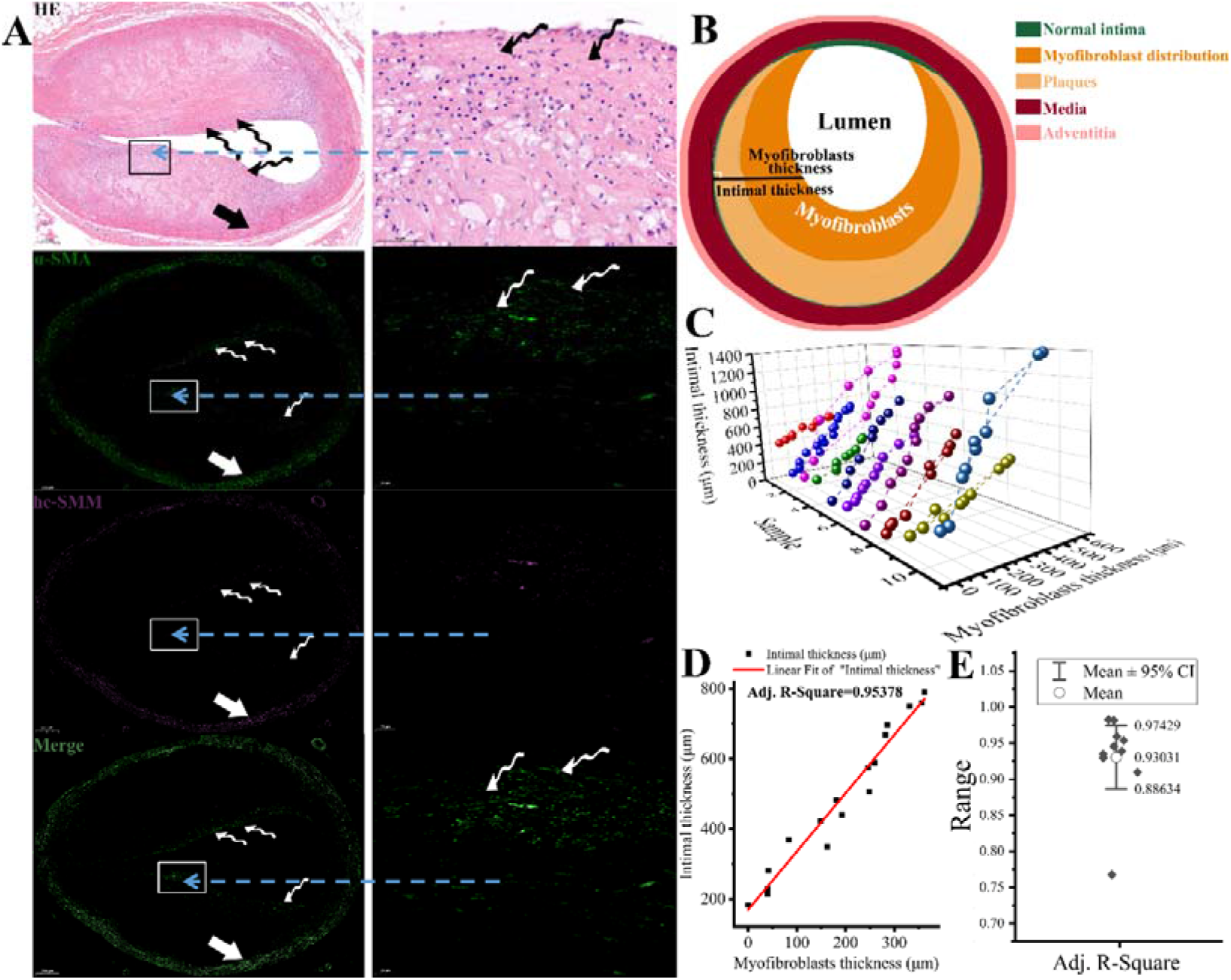
Myofibroblasts in human coronary atherosclerotic plaques. **(A)** It was the HE and immunofluorescence staining of the same coronary artery. HE staining showed that the intimal thickness and atherosclerotic degree were different in different positions of the same transection. There were more cells in the intima near the lumen in the more severe plaque areas. Immunofluorescence staining showed that there were more cells (curved arrows) highly express α-SMA but lowly express hc-SMM in the intima near the lumen with more severe atherosclerosis. They were myofibroblasts rather than SMCs (straight arrows) which highly express both α-SMA and hc-SMM in the media of arteries. **(B)** It is the pattern diagram of myofibroblasts distributed in atherosclerotic plaques. **(C)** There were no myofibroblasts distribution in the arteries without atherosclerosis. To study the relationships between atherosclerotic plaques and myofibroblasts, the arteries with obvious atherosclerosis were analyzed here. 10-20 different locations were randomly selected from the same transection of each coronary artery. The myofibroblasts thickness and intimal thickness were measured carefully in each artery transection. Arteries with obvious atherosclerosis(n=10) were analyzed and got the scatter diagram of myofibroblasts thickness and intimal thickness. **(D)** Linear fit between them was done and got the Adj. R^2^ in each artery transection. **(E)**These Adj. R^2^ were analyzed (e, 95% confidence interval was 0.89 to 0.97). Myofibroblasts thickness is highly positive correlated with intimal thickness. It is myofibroblasts play an important role in the formation of atherosclerotic plaques.

### Collagen fibers were the main components of human coronary atherosclerotic plaques

Collagen fibers were forming in the intima near the lumen (fibrous cap), while the collagen fibers far from the lumen were denaturing, and some of them kept the original structure and some of them had lost their original structure (Fig.2A, 3). In the control (Con) group, the proportion of collagen fibers in intima was low, while in the atherosclerosis (AS) group, it was significantly bigger (Fig.2B, 2C, P<0.01). It showed that when the ratio of collagen fibers was less than 30%, the intima thickening was not obvious, while the ratio was more than 40%, the intima began to thicken obviously. Of course, due to the difference of artery diameter, this threshold is not a definite value. It is a range, which is about 30%-40% in human coronary arteries. The formation and degeneration of fibrous tissues would lead to formation of fibrous plaques and various composite plaques (Fig.2, 3). The distribution of the newborn collagen fibers coincided with that of myofibroblast (Fig.1), both in the intima near the lumen (fibrous cap).

**Fig.2.**
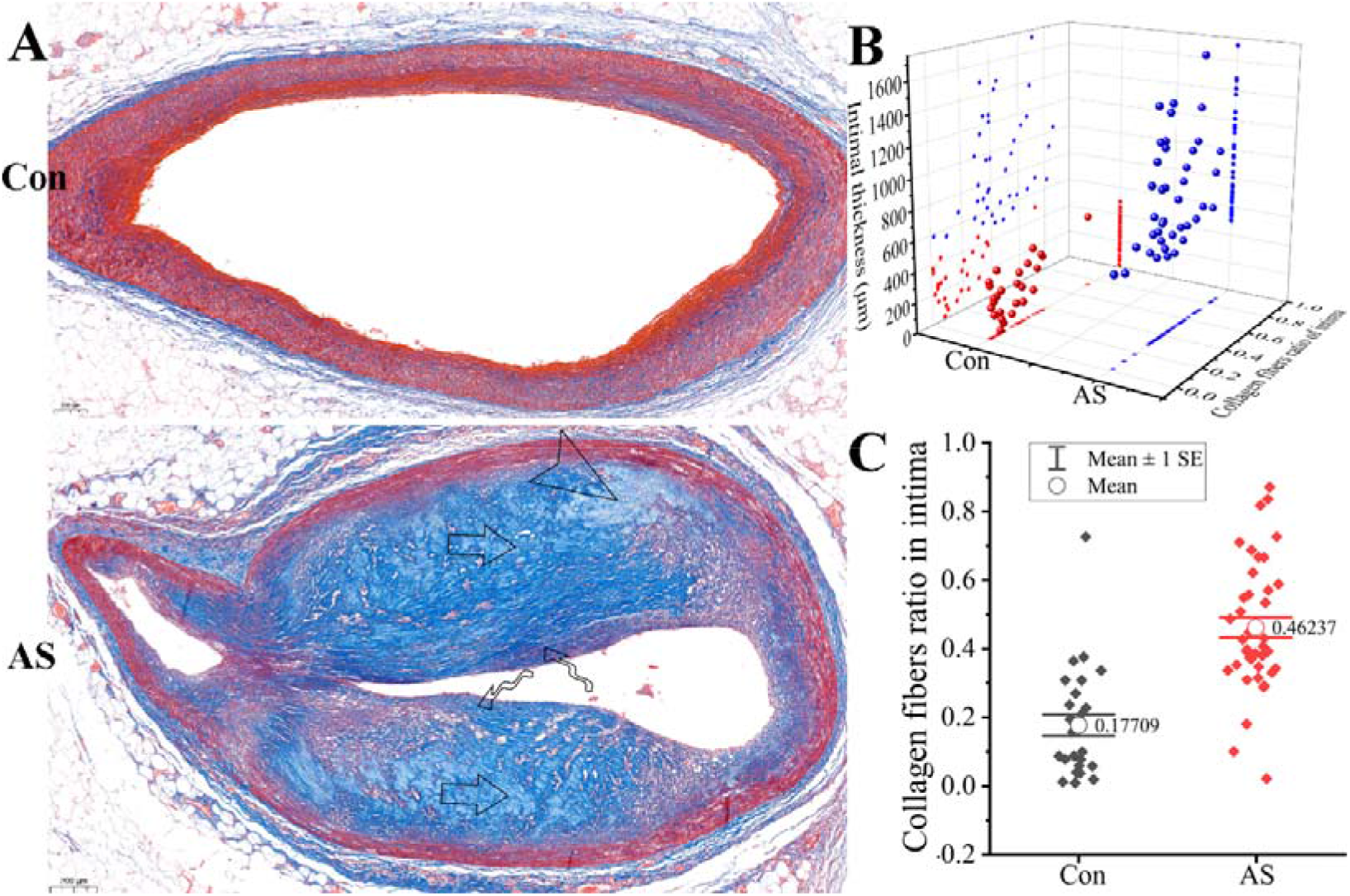
Collagen fibers of intima from human coronary arteries. **(A)** It was the Masson staining of human coronary arteries and collagen fibers were blue stained. There is no clear definition of atherosclerosis. In our study, for statistical convenience, we defined atherosclerosis of coronary artery as intimal thickness (the thickest location of intima) > 500μm and the samples were divided into control (Con) group and atherosclerosis (AS) group. The collagen fibers were few in Con group (A, Con, n=26), While collagen fibers were widely distributed in AS group (A, AS, n=40). Collagen fibers were forming in the intima near the lumen (newborn, curved arrows), while the collagen fibers far from the lumen were denaturing, and some of them kept the original structure (straight arrows) and some of them had lost their original structure(arrowheads). (**B**) It was a scatter diagram of the ratio of collagen fibers in the intima and the intimal thickness (the thickest location of intima). (**C**)There was significant difference in the ratio of collagen fibers between the two groups with T test (P<0.01).

**Fig. 3.**
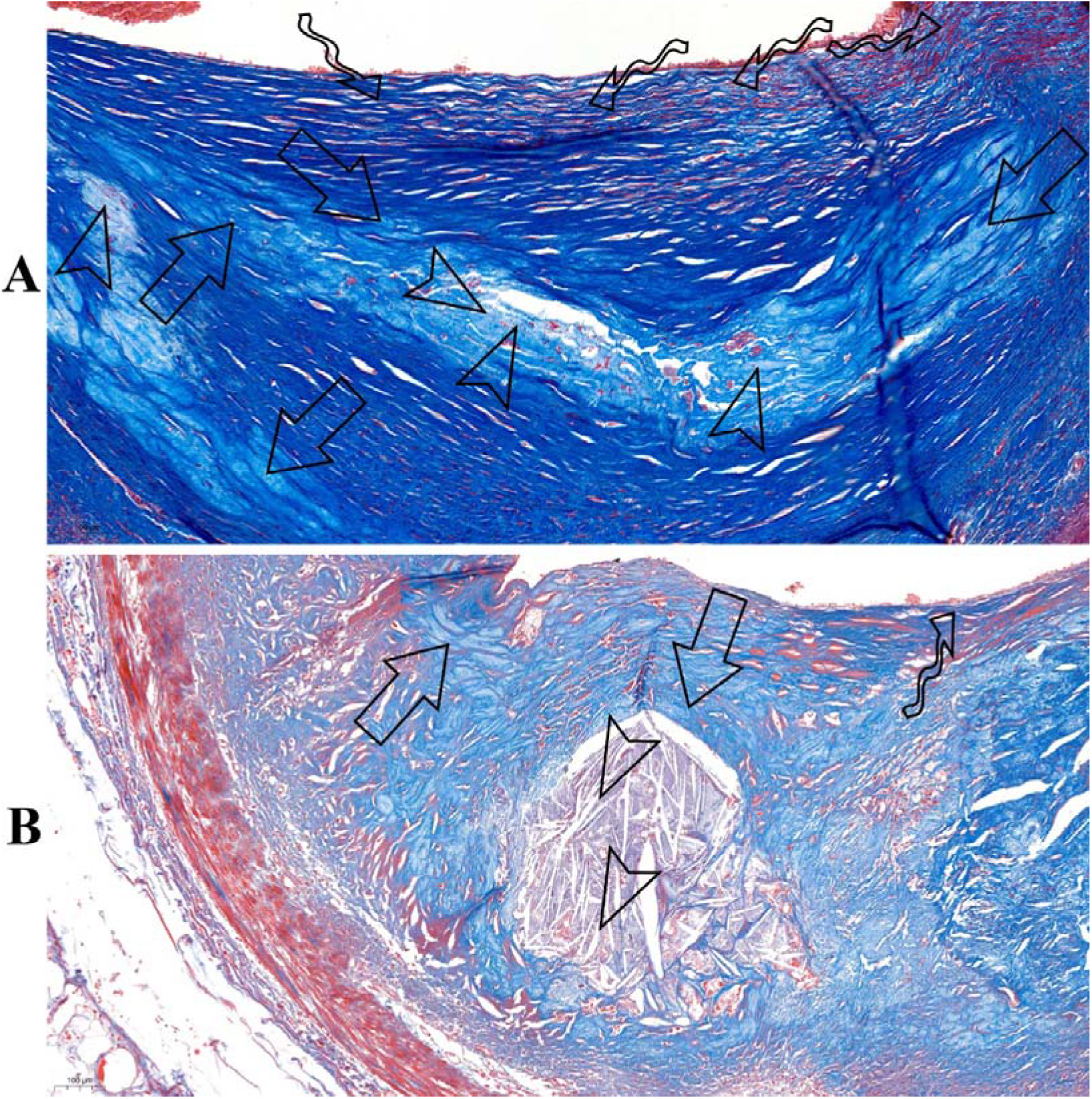
Formation of soft plaques or calcified plaques in human coronary arteries. Collagen fibers were stained blue with Masson staining. (**A**) The new born collagen fibers were forming in the intima near the lumen (fibrous cap, curved arrows). Some collagen fibers far from the lumen had denatured dyed with light blue, lost their normal structure and support capability (arrowheads), and soft plaques is gradually forming. Around the denatured collagen fibers (arrowheads), there were denaturing collagen fibers (straight arrows) kept the relatively normal structure of fibers. (**B**) The denaturation of collagen fibers is much more serious, and most of them had denatured. There were crystalline calcified tissues (arrowheads) filled parts of the denatured collagen fibers and calcified plaques had formed here. These phenomena are not individual cases, but common in human coronary atherosclerotic plaques.

### Myofibroblast formed lipid rich plaques with foam cells, extracellular lipids and collagen fibers in vitro

In order to verify its function, myofibroblasts were cultured in vitro. In this study, we did not add modified low-density lipoprotein (LDL) and just used 5% FBS high glucose DMEM which contained low amounts of native LDL as the culture medium. LDL could be taken by myofibroblasts and promote cell proliferation (Fig.4A, 4B). Collagen fibers containing lots of lipids (Fig.5A) and foam cells widely formed in vitro (Fig.4C, 4D, Fig.5). Extracellular lipids accumulation and lipid rich plaques could also be perfectly presented (Fig.4C, 4D, Fig.5). Myofibroblasts derived foam cells could form in or around the lipid rich plaques in large quantities, which looked like soap bubbles and were full of lipid droplets accumulation in the cytoplasm (Fig.5B, 5C). And intracellular lipids leaked out of cells, resulting in the accumulation of extracellular lipids in the plaques (Fig.4C, 4D, Fig.5).

**Fig.4.**
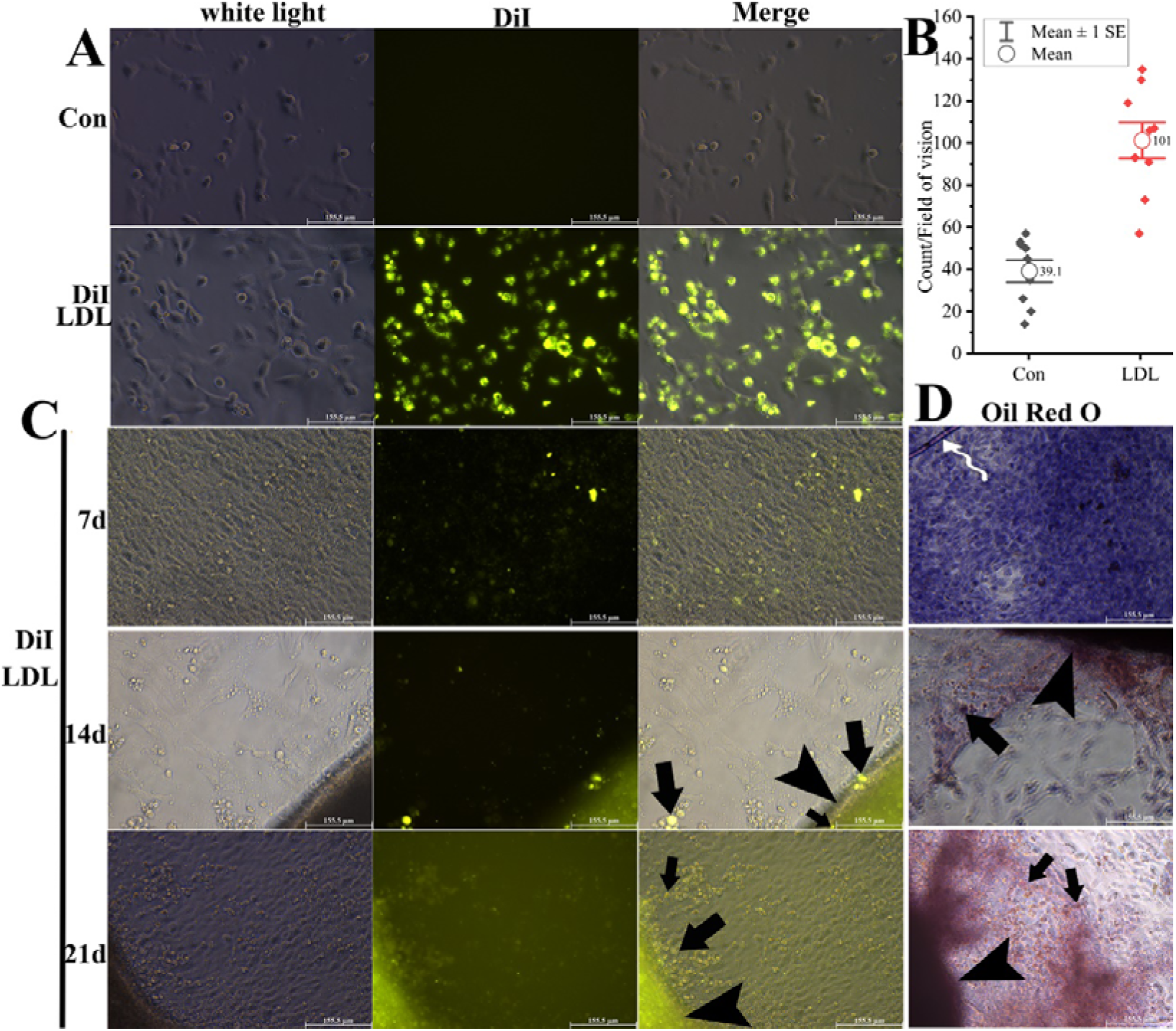
Myofibroblasts in vitro formed atherosclerotic plaques with lots of foam cells and extracellular lipids. **(A)** LDL could be heavily utilized by myofibroblasts which were labeled with strong fluorescence of DiI in DiI-LDL group at 24 hours. **(B)** Cell proliferation in DiI-LDL group was much faster than that of the control group at 24 hours, which was analyzed with T test (Con, b, n=9, P<0.01). Although the cells took lots of LDL, the cell morphology at this stage had not changed significantly and could not be called foam cells. **(C)** The myofibroblasts labeled with DiI-LDL continue to be cultured for 7, 14 and 21 days in 5% FBS high glucose DMEM with 5% CO_2_ at 37◻ and be imaged. **(D)** Oil red O staining for lipids was performed at 7 days, 14 days or 21 days respectively. Cells covered the entire well and DiI fluorescence was dispersed throughout the well due to cell proliferation at 7 days. During this period no obvious foam cells formed. At 14 days and 21 days, myofibroblast derived foam cells round shaped with strong DiI fluorescence (C, straight arrows) and dyed red with Oil red O staining for lipids (D, straight arrows) were widely distributed in or around the plaques. Lipid rich plaques with strong DiI fluorescence (C, arrowheads) and dyed red with Oil red O for lipids (D, arrowheads) widely formed at the time. We labeled myofibroblasts with DiI-LDL(A). Now the lipid rich plaques with foam cells (C, straight arrows) and lipids (C, arrowheads) all showed high intensity fluorescence of DiI. This proved the leakage of intracellular lipids to extracellular forming the extracellular lipids in the plaques (C, D, arrowheads).

**Fig.5.**
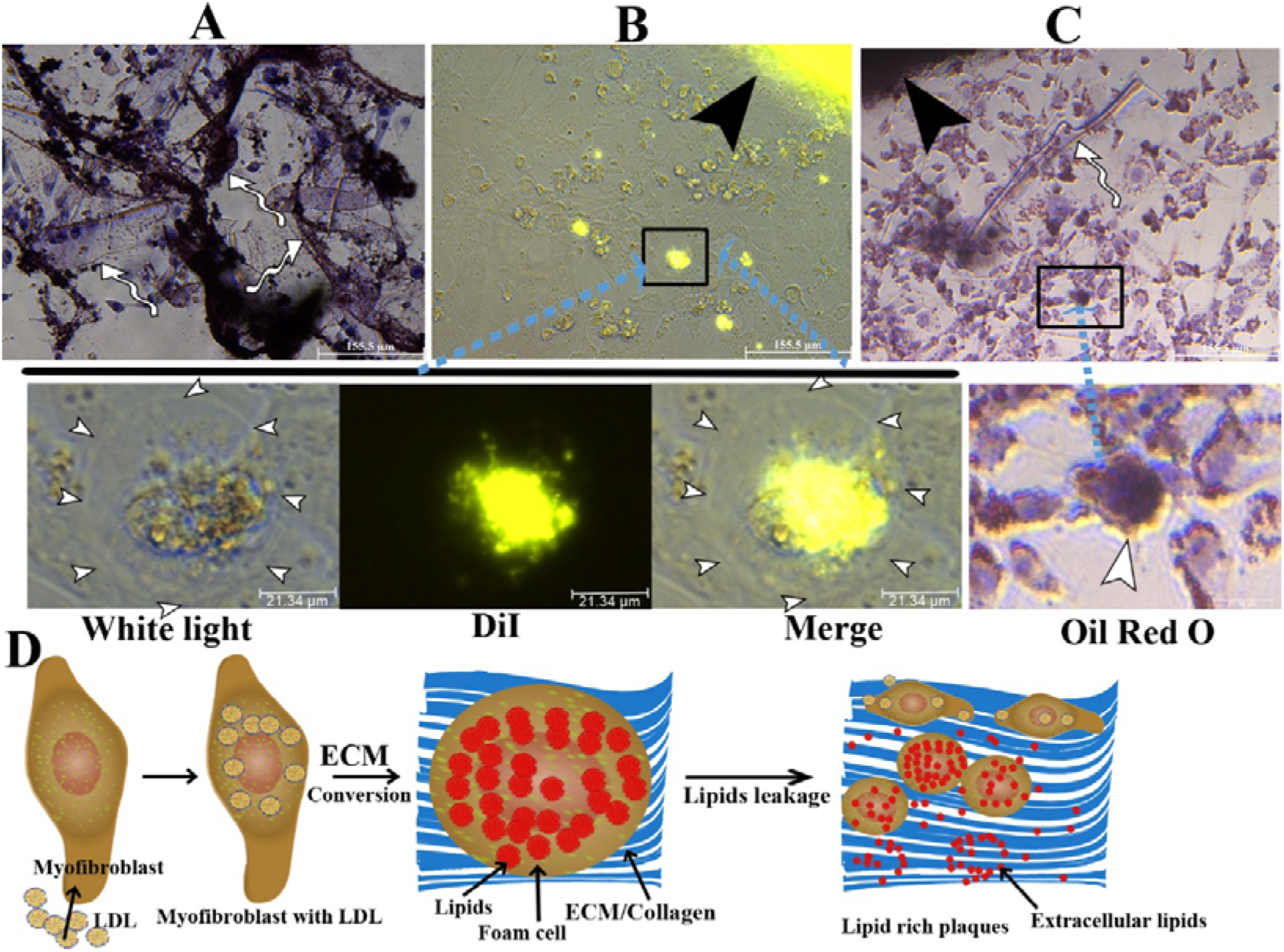
Myofibroblasts formed lipid rich plaques with collagen fibers, foam cells and extracellular lipids. Myofibroblasts were cultured in vitro at days 14. (**A**) Extracellular matrix (ECM)/Collagen fibers formed widely, and contained lots of lipids which could be deeply dyed by Oil Red O (curved arrows, n=9). Myofibroblasts derived foam cells (**B, C**, white arrowheads showing cell membrane) in or around the lipid rich plaques (**B, C**, black arrowheads). Foam cell was round with high intensity fluorescence of DiI (**B**) and full of lipids(**C**). **(D)** It illustrated the process of myofibroblast forming lipid rich plaques with collagen fibers, foam cells and extracellular lipids. Myofibroblasts are like LDL processing factories in the intima near the lumen, which could take LDL to promote proliferation, and convert the proteins part of lipoproteins into extracellular matrix (ECM) and exocytosis outside forming collagen fibers, while the lipids part of lipoproteins is deposited in the cell forming lipid droplets in foam cells. Lipids in the cells leaked out forming extracellular lipids accumulation.

## Discussion

Hypertension, diabetes, hyperlipidemia, smoking or vasculitis, etc. are the common risk factors of atherosclerosis. These risk factors could damage blood vessels through physical, chemical or biological ways. The large and medium arteries are responsible for transporting flowing blood from the heart to organs of the body. They must bear much bigger blood pressure and the flowing blood. If there is a serious damage in one position and lack effective tissue repair, it would lead to aneurysms, dissections, or even rupture of large and medium arteries^18^, which would be fatal to the body. Fortunately, vascular tissue repair, characterized by fibrous tissue remodeling, could alleviate these injuries, so the incidence of these acute arterial rupture diseases is not so high^19^. However, vascular fibrous tissue remodeling is a double-edged sword. If it could not be well regulated, it would lead to excessive proliferation of fibrous tissues, degeneration and necrosis of them, which would lead to the formation of fibrous plaques or composite plaques. In this study, collagen fibers were forming in the thickening intima near the lumen (fibrous cap) leading to fibrous plaques, and were denaturing or calcifying far from the lumen evolving into various complex plaques (Fig.2, 3).

In vascular fibrous repair and remodeling, which cell take primary responsibility? To clarify this question, we observed human coronary arteries and found that in atherosclerotic plaques, SMCs were few, while lots of myofibroblasts infiltrated, especially in the intima near the lumen. what’s more, their distribution was highly positive related to intimal thickness (Fig.1). It suggests that myofibroblast play a major role in human coronary atherosclerotic plaques formation. Although myofibroblasts are similar to SMCs in morphology, they are different in cell proliferation and extracellular matrix secretion^10, 16^. α-SMA is highly expressed in both SMCs and myofibroblasts, while hc-SMM is highly expressed in SMCs but lowly expressed in myofibroblasts^9, 17^. Many previous studies used the high expression of α-SMA to identify SMCs, which might lead to myofibroblasts being mistaken for SMCs. Moreover, SMCs are stable cells, and their ability of regeneration and repair is weak^10^. Unlike SMCs, myofibroblast is widely distributed in the skin, liver, lung, etc.^9^, especially in the repair stage of organ injury^16^. Its strong ability of proliferation and secretion of extracellular matrix provides support for its repair and remodeling of damaged vessels. In our study, the position of collagen fibers generated (fibrous cap) were consistent with that of myofibroblasts (Fig.1).

Because of their strong ability to proliferate and secrete extracellular matrix, myofibroblasts need lots of nutrients. LDL has a diameter of 20 nm, and an average composition of 20% proteins, 20% phospholipids, 40% cholesteryl esters (CEs), 10% unesterified cholesterol (UC), and 5% triglycerides (TGs)^20^, which could provide large numbers of lipids and proteins for myofibroblasts. In vitro, LDL could be taken by myofibroblasts and promote cell proliferation (Fig.4). Foam cells are characterized by lots of lipid droplets accumulation in the cytoplasm and certain morphological changes^13^. Previous studies in vitro showed that native LDL could not cause excessive intracellular lipids accumulation and the consequent foam cells formation on SMCs or macrophages^13, 21^. Therefore, the most popular method to do that now is using modified LDL, such as oxidized LDL (ox-LDL), to establish SMCs or macrophage derived foam cells^22, 23^. However, foam cells formed in this way were far from those in human atherosclerotic plaques, for example, SMCs did not show obvious roundness. Unlike previous studies, we did not add modified LDL and just used 5% FBS high glucose DMEM which contained low amounts of native LDL as the culture medium, myofibroblasts in vitro widely formed lots of foam cells, and collagen fibers and extracellular lipids could also be perfectly presented. According to our results (Fig.5), myofibroblasts derived foam cells could form in or around the lipid rich plaques in large quantities, which looked like soap bubbles and were full of lipid droplets in the cytoplasm, and lots of collagen fibers contained extracellular lipids also formed. It is well known that myofibroblasts could secrete large numbers of extracellular matrix forming collagen fibers. Meanwhile, in vitro without other interfering factors, we believe that these extracellular lipids were derived from the exocytosis of intracellular lipids. In conclusion, lipid rich plaques are composed of extracellular matrix/collagen fibers, foam cells and extracellular lipids. This is consistent with the characteristics of human coronary atherosclerotic plaques. This method of lipid rich plaques formation in vitro is simple, easy to repeat, and does not need the intervention of other factors such as modified LDL, which would provide a new and high credibility assistance for the future study of atherosclerosis. We also cultured macrophages or SMCs respectively in the same conditions, but did not form lipid rich plaques. This suggests that myofibroblasts should be the direct source of atherosclerotic plaques. It would provide a target cell for future treatments of atherosclerotic diseases.

Human myofibroblasts in vitro formed lipid rich plaques in normal culture conditions without other intervention. It suggests that in addition to the formation of fibrous tissues, myofibroblasts could also play an important role in foam cells formation and extracellular lipids accumulation. Benditt EP proposed the hypothesis of atherosclerosis caused by the “mutant SMC”, which also mentioned “SMC transformation”^14, 15^. Our findings not only do not conflict with this theory, but further develop it, making the pathogenesis of atherosclerosis clearer. Myofibroblast is not found in normal blood vessels in our study, and it might be transformed from other cells in pathological conditions. In the injured skin, lung or liver, etc., myofibroblast could be transformed from fibroblasts, endothelial cells or SMCs^9^. In blood vessels, it is still unclear what kind of cells myofibroblast comes from, and it might transform from SMCs, fibroblasts, endothelial cells, etc., which are the common vascular resident cells. Therefore, we speculate that myofibroblast might be derived from SMC, and the transformation from fibroblast or endothelial cell could not be excluded, which still needs to be clarified in future studies. The traditional concept is that the pathogenesis of atherosclerosis is very complicated. Serious tissue damage or bigger mechanical force increase could promote myofibroblasts transformation from other cells^16, 24^. Hyperglycemia or hyperlipidemia could provide much more glucoses, proteins and lipids for myofibroblasts. These risk factors are the promoting factors of atherosclerosis by damaging blood vessels, or by promoting transformation or proliferation of myofibroblasts. Macrophages are involved in fibrous tissue remodeling, are trapped in atherosclerotic plaques and form macrophage derived foam cells. We studied the roles of myofibroblast in atherogenesis at the cellular level. Lots of molecular studies need to be further explored in the future.

## Conclusions

Atherogenesis should be mainly a process of myofibroblast repair and remodeling. It is myofibroblasts which are closely related to collagen fibers formation and intimal thickening. LDL could be taken by myofibroblasts to enhance proliferation. It’s the first time that lipid rich plaques formation was presented with human myofibroblasts cultured in vitro. It suggested that myofibroblast should be the direct and main performer of atherosclerotic plaques formation. It would provide a target cell for future treatments of atherosclerotic diseases. In vitro atherosclerotic plaques model formed by myofibroblasts would provide an efficient and convenient way to study atherosclerosis.

## Acknowledgements

This work was supported by China Postdoctoral Science Foundation (no. 2018M641934) and a grant of Innovative Research Groups of the National Natural Science Foundation of China (81521001). Thanks for Zhengdong Li from Institute of Forensic Sciences of Shanghai providing the autopsy specimens.

## Author contributions

Junbo Ge supervised the study. Junbo Ge and Xinggang Wang designed the experiments. Xinggang Wang performed experiments, analyzed data, and wrote the manuscript. Junbo Ge made manuscript revisions.

